# Spatial transcriptomics of healthy and fibrotic human liver at single-cell resolution

**DOI:** 10.1101/2024.02.02.578633

**Authors:** Brianna Watson, Biplab Paul, Liat Amir-Zilberstein, Asa Segerstolpe, Raza Ur Rahman, Angela Shih, Jacques Deguine, Ramnik J. Xavier, Jeffrey R. Moffitt, Alan C. Mullen

## Abstract

Single-cell RNA sequencing (scRNA-seq) has advanced our understanding of cell types and their heterogeneity within the human liver, but the spatial organization at single-cell resolution has not yet been described. Here we apply multiplexed error robust fluorescent in situ hybridization (MERFISH) to map the zonal distribution of hepatocytes, resolve subsets of macrophage and mesenchymal populations, and investigate the relationship between hepatocyte ploidy and gene expression within the healthy human liver. We next integrated spatial information from MERFISH with the more complete transcriptome produced by single-nucleus RNA sequencing (snRNA-seq), revealing zonally enriched receptor-ligand interactions. Finally, analysis of fibrotic liver samples identified two hepatocyte populations that are not restricted to zonal distribution and expand with injury. Together these spatial maps of the healthy and fibrotic liver provide a deeper understanding of the cellular and spatial remodeling that drives disease which, in turn, could provide new avenues for intervention and further study.

## Introduction

The liver is composed of parenchymal, non-parenchymal, and immune cells that are organized into anatomic structures called lobules, which are 0.5-1 mm in diameter and defined by sites of portal venous and arterial inflow and central venous outflow^1, 2^. Within the lobule, hepatocytes are classically organized into three zones, with zone 1 proximal to the portal vein and hepatic artery, zone 3 proximal to the central vein, and zone 2 in the intermediate region^3,4^. Hepatocyte zonation matches function to the physiologic environment. For example, hepatocytes in zone 1 are more involved in gluconeogenesis and beta-oxidation, reflecting the relatively oxygen and nutrient rich region near the portal vein and hepatic artery, whereas hepatocytes in zone 3 are more active in glycolysis and lipogenesis, reflecting the depletion of oxygen and nutrients near the central vein^3–5^. In this sense, such zonal organization is critical for our understanding of hepatocyte activity. However, the degree to which hepatocytes adjust their gene expression and, thus, function within the spatial context in the liver remains unclear, as current single-cell analysis of the human liver have either measured gene expression without spatial data or evaluated spatial data without single-cell resolution.

The liver is also composed of non-parenchymal cells, which include hepatic stellate cells (HSCs), liver sinusoidal endothelial cells (LSECs), resident macrophages, and cholangiocytes^6,7^. While sub-populations of macrophages and HSCs in the healthy human liver have been described in single-cell data^8–10^, these subpopulations, and potential gene expression variations within, have not been resolved within the lobule organization at single-cell resolution in the human liver. As such, their role within the spatial context of the liver remains unclear.

In addition to spatial location, another feature of hepatocytes that may drive functional heterogeneity is their ploidy^11^. Specifically, hepatocytes in the adult liver can either be single or multinucleated, and nuclei themselves can vary in ploidy. Early in life, most human hepatocytes are diploid and contain a single nucleus; ploidy increases with age such that in adults, about a third of hepatocytes are multinucleated, or their nuclei contain more than two copies of each chromosome^12–14^. Yet the spatial organization of multinucleated hepatocytes and the impact of nuclear content on gene expression remains poorly understood^15^.

Here, we apply image-based spatial transcriptomics (multiplexed error robust fluorescence in situ hybridization; MERFISH) and single-nucleus RNA sequencing (snRNA-seq) to the same samples of healthy human liver in order to construct spatial maps of hepatocytes and non-parenchymal cells at single-cell and transcriptome-wide resolution. This analysis allowed us to define gradients of gene expression across human hepatocyte zones, determine the cell types where receptor-ligand co-expression are in spatial proximity to promote crosstalk, and evaluate the relationship between ploidy and gene expression within hepatocytes. We performed similar measurements in fibrotic liver samples to understand the changes in hepatocytes and non-parenchymal cells that occur with chronic injury. These analyses provide a single-cell spatial transcriptomic map of the human liver, annotating the gradient in hepatocyte gene expression from the portal area to central vein and show that hepatocyte ploidy is equally distributed across zones within the lobule and does not affect differential gene expression. In addition, we define subsets of spatially distinct macrophages and HSC populations, and identify the new expansion hepatocyte populations with chronic injury, which together provide an approach and framework to understand changes that occur with human liver disease at single-cell resolution in space.

## Results

### Spatial organization of hepatocytes in the human liver

To explore the spatial and cellular organization of the human liver, we collected human liver tissue from surgical resections of three adult patients (two female and one male; **Supplementary Table 1**). We designed a MERFISH panel targeting 317 genes with a focus on hepatocytes (**Methods**), and then characterized the expression of these genes within cryosections taken from these samples using MERFISH^16–18^.

The distribution of key individual genes revealed the rich spatial architecture of the liver and provided an initial validation of our measurements. We observed sheets of cells expressing *ALDOB*, consistent with hepatocytes (**Supplementary Fig. 1**), while *CD5L* expression was clustered in spaces between hepatocytes consistent with resident macrophages located in the sinusoids (**Supplementary Fig. 1a-f**). *PDGFRB* expression was concentrated along the edges of ALDOB-expressing hepatocytes in a distribution expected for HSCs, which are located in the subendothelial space (**Supplementary Fig. 1a-f**). *KRT7* and *SOX9* were enriched in periportal areas, often in larger clusters consistent with cholangiocytes (**Supplementary Fig. 1g-l**), while *DNASE13* and *INMT* expression were scattered through the parenchyma consistent with the distribution of LSECs (**Supplementary Fig. 1g-l**).

To define individual cells within our data, we included in our MERFISH measurements an immunofluorescence stain against a pan-cell surface marker, the Na^+^/K^+^-ATPase^19^. This antibody stain was incorporated into our MERFISH readout through the use of an oligonucleotide-tagged secondary antibody^19^ (**Methods**). We then leveraged Cellpose^20^ to define cell boundaries in three dimensions from these co-stains, assigned RNAs within these boundaries, and then used an RNA-based segmentation routine, Baysor^19^, to adjust and improve these boundaries and recover cells for which the co-stains did not provide clear boundaries. Following this analysis and cuts for cells with small numbers of transcripts, we compiled a dataset of ∼110,000 cells from the healthy human liver.

The expression of all 317 genes was quantified in each cell to generate a count matrix for the human liver samples. The count matrix for the pooled data was then visualized by Uniform Manifold Approximation and Projection (UMAP) after the application of batch correction (**Methods; Fig. 1a**). Three clusters of hepatocytes were defined along with two HSC clusters, two macrophage clusters, one LSEC cluster, and one cholangiocyte cluster. Each of these clusters were defined by classic markers, supporting our cluster assignment (**Fig. 1b**). We noted that additional immune cell populations were not resolved in this analysis, likely reflecting the choice of genes included in our panel.

**Figure 1.**
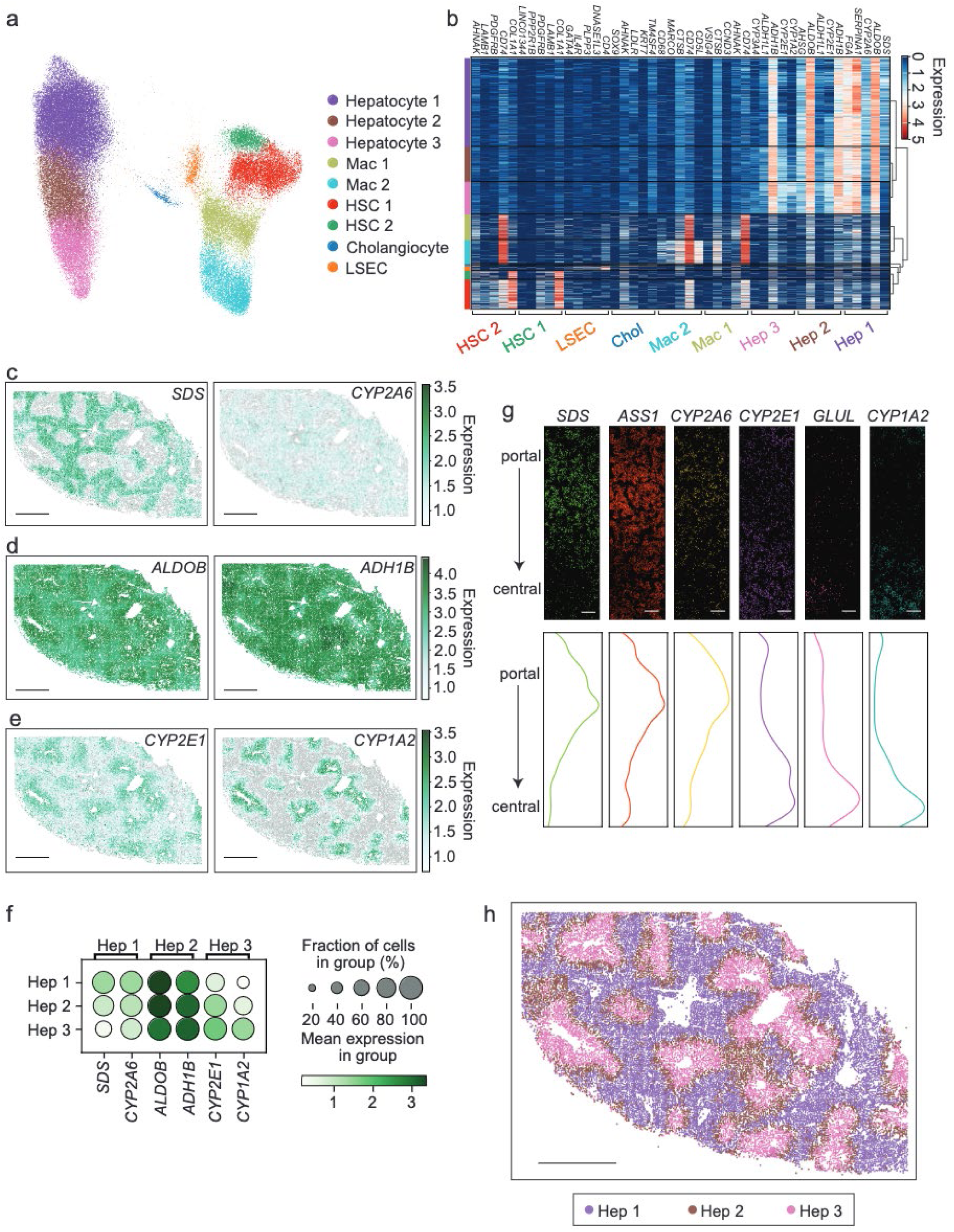
Mapping hepatocytes within the architecture of the healthy human liver with MERFISH. **a.** UMAP of all cells measured with MERFISH in the healthy human liver. **b.** Heatmap displaying differentially expressed genes that identify each cluster. Normalized gene expression is indicated by color. Each column represents a single cell, and cells are grouped by cluster as indicated by color at the left of the heatmap. Genes are indicated on the top of the heatmap, and groups of genes characteristics of individual clusters are indicated on the bottom. Hierarchical clustering is shown at the right. Genes enriched in each cluster are shown. Genes may appear more than once if enriched in more than one cluster. **c-e.** Distribution of cells expressing *SDS* and *CYP2A6* (enriched in zone 1, **c**), *ALDOB* and *ADH1B* (most enriched in zone 2, **d**), and *CYP2E1* and *CYP1A2* (enriched in zone 3, **e**) in a section of liver tissue. Scale bars: 1000 μm. Expression level is measured in normalized gene expression. **f**. Dot plot quantifying expression of genes mapped in **c-e** where the size of the dot represents the percentage of cells expressing a specific gene, and the color intensity indicates mean expression. **g.** mRNA distributions across a single lobule. Each dot represents an individual mRNA transcript for the indicated gene oriented from a portal region at the top to a central region at the bottom. The relative abundance of each transcript across the lobule was then plotted (lower panel) for the same field of view. Scale bars: 50 μm. **h.** Spatial distribution of cells assigned to each cluster of hepatocytes (Hep 1, Hep 2, Hep 3). Hep 1 cells (purple) map to periportal areas (zone 1), Hep 3 cells (pink) map to pericentral areas (zone 3), and Hep 2 cells (brown) map between Hep 1 and Hep 3 cells (zone 2). Scale bar: 1000 μm.

We next explored the spatial distribution of hepatocytes by evaluating the expression of individual genes (**Fig. 1c-f**, **Supplementary Fig. 2a**). *SDS* was enriched in zone 1 (periportal), while *CYP2E1* expression increased in zone 3 (pericentral), as previously described in mouse^21–24^ (**Fig. 1c, e, f; Supplementary Fig. 2a**). In addition, we found that *CYP2A6* and *ASS1* were enriched in portal regions similar to *SDS,* and *CYP1A2 and GLUL* were enriched in central regions similar to *CYP2E1 (***Fig. 1c, e, f-g**). *ALDOB* and *ADH1B* were expressed in all hepatocytes but were enriched in zone 2, with *ALDOB* expression shifted towards zones 1 and 2 and *ADH1B* expression shifted towards zones 2 and 3 (**Fig. 1d, f**). These results showed that MERFISH captures differences in gene expression across zones within the hepatic lobule, and we next asked how effectively we could define zonation with the full MERFISH probe set. Mapping hepatocyte clusters 1, 2, and 3 (**Fig. 1a**) back to tissue sections showed clear zonal distribution, with hepatocyte cluster 1 localizing to zone 1, hepatocyte cluster 3 to zone 3, and hepatocyte cluster 2 to zone 2 (between zone 1 and 3; **Fig. 1h, Supplementary Fig. 2b, c**). Collectively, these observations are consistent with the classic zonation described in human and mouse liver^4,21–23^, supporting our MERFISH measurements.

While our analysis revealed that hepatocytes can be separated into three populations, we noted that these populations are contained within a single group of cells in the UMAP representation of the data (**Fig. 1a**). This representation suggests that the classic liver zones represent approximations of a continuous variable in gene expression in hepatocytes determined by the spatial relationship between portal and central areas of the lobule. To explore this possibility further, we performed a pseudotime analysis on gene expression of hepatocytes (**Supplementary Fig. 2d)** and found that the pseudotime mapped to a continuous distribution in space (**Supplementary Fig. 2d-f**). This gradient of gene expression could also be observed across individual lobules (**Fig. 1g**), and many genes show a gradient in gene expression from periportal to pericentral regions (**Supplementary Fig. 2g**). Collectively, these observations suggest that spatial gene expression in human hepatocytes is better described by a continuous variable determined by the relative distance between the portal and central areas of the lobule, similar to observations in mouse^23,24^.

### Spatial organization of non-parenchymal cells in the human liver

We next expanded our analysis to include the spatial organization of non-parenchymal cell types (**Fig. 2a, b**). Two macrophage populations were visualized in the healthy liver and showed different patterns of distribution (**Fig. 2c, Supplementary Fig. 3a**). Macrophage (Mac) cluster 1 was enriched in the periportal area and also dispersed through the lobules, while cells from Mac 2 were scattered more diffusely through the lobules. Both macrophage populations were identified by expression of *CD74*, while Mac 2 cells also expressed *CD5L* and *MARCO* (**Fig. 2d, i; Supplementary Fig. 3b)**, most consistent with non-inflammatory macrophages or Kupffer cells^8,9,25^.

**Figure 2.**
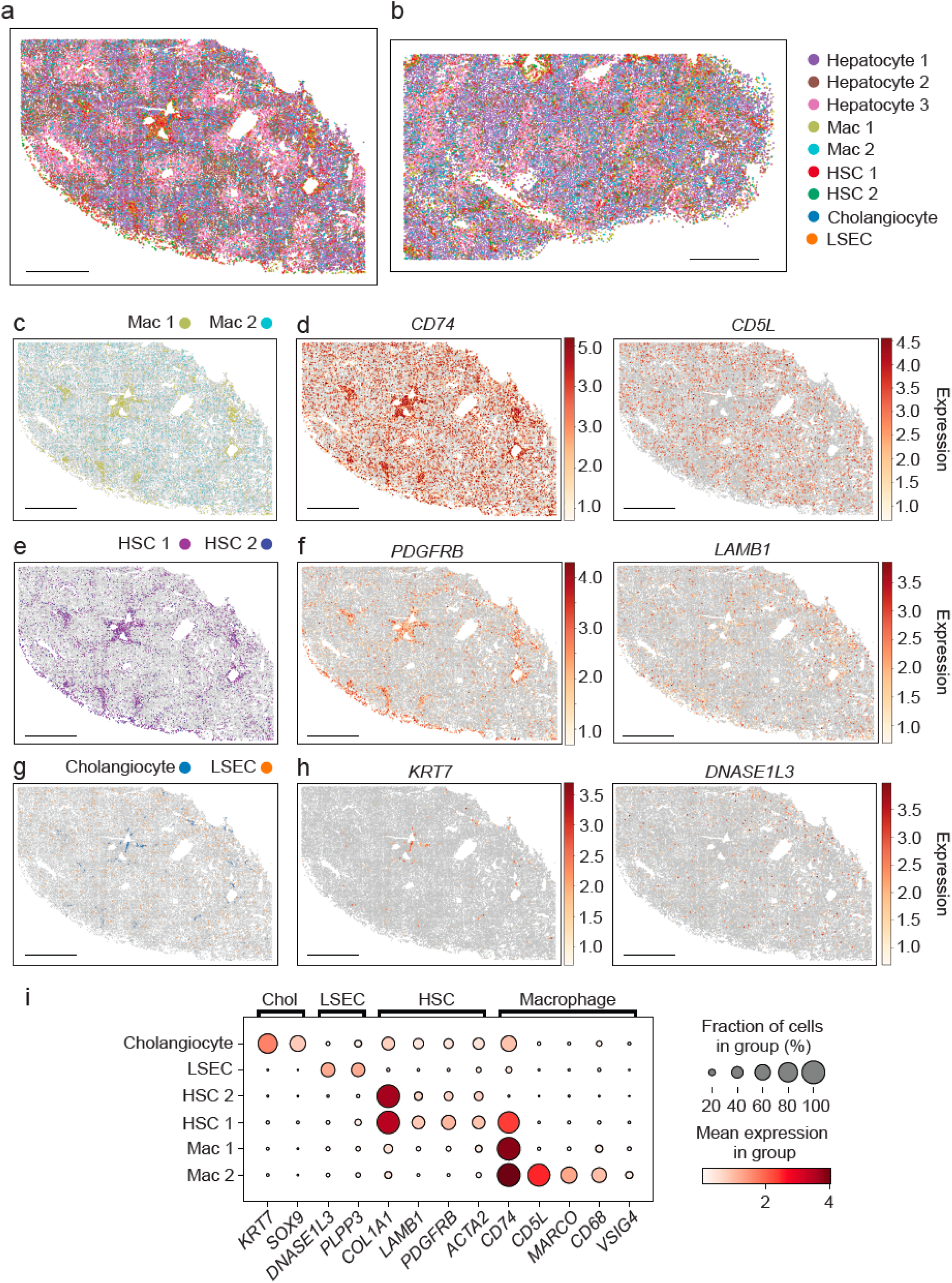
MERFISH maps non-parenchymal cells within the healthy liver. **a,b.** Spatial distribution of all cells defined by MERFISH within the architecture of the healthy liver for tissue from two different donors, donor 1 (**a**) and donor 2 (**b**). Scale bars: 1000 μm. **c,e,g**. Spatial distribution of macrophage (**c**), HSC (**e**), and cholangiocyte and LSEC (**g**) populations mapped in the same tissue section as **a**. Scale bars: 1000 μm. **d, f, h**. Spatial distribution of cells in the same tissue section as (**a**) colored by the normalized expression of the indicated marker genes. *CD74* and *CD5L* identify macrophages (**d**), *PDGFRB* and *LAMB1* identify HSCs (**f**), KRT7 identifies cholangiocytes, and *DNAS1L3* identifies LSECs (**h**). Scale bars: 1000 μm. **i.** Dot plot quantifying expression of genes that identify non-parenchymal cells in the liver. The size of the dot represents the percentage of cells expressing a specific gene, and the color intensity indicates mean expression.

Two clusters were also identified demonstrating patterns of gene expression consistent with HSCs (**Fig. 2e; Supplementary Fig. 3c**). These cells also displayed different spatial distributions, with cells from HSC 1 enriched in periportal regions and spread through the lobules, while cells from HSC 2 were scattered more diffusely through the lobules without periportal enrichment. These two populations differed primarily in the levels of *CD74* expression, while both populations expressed *PDGFRB* and *LAMB* (**Fig. 2f, i; Supplementary Fig. 3d**).

Cholangiocyte and LSECs were also identified by MERFISH and followed predicted distributions (**Fig. 2g; Supplementary Fig. 3e**). Cholangiocytes were localized to portal regions and were identified by expression of *KRT7* (**Fig. 2h, left; Supplementary Fig. 3f, left)**, while LSECs, which line the sinusoids, were identified more diffusely through the liver, and expressed *DNASE1L3* (**Fig. 2h, right; Supplementary Fig. 3f, right**). We did not note clear patterns of zonation in gene expression within these non-parenchymal populations^26–28^, suggesting that, if such gradients exist in the human liver, our MERFISH library does not contain the proper genes to capture them.

### Combining MERFISH with snRNA-seq to integrate the full transcriptome in space

MERFISH identifies parenchymal and non-parenchymal cells in human liver tissue at single-cell resolution and provides expression data for hundreds of genes, but it does not cover the full transcriptome. To address this limitation, we next performed snRNA-seq analysis for the same human liver samples analyzed by MERFISH, profiling, after filtering, ∼15,000 nuclei. To combine these different measurement modalities, we leveraged standard data integration tools (**Methods**) and then jointly clustered the integrated data (**Supplementary Fig. 4a**).

As expected, this joint analysis revealed a similar diversity of cell populations in the snRNA-seq data to what we observed in the MERFISH data (**Fig. 3a, b**). We identified three major populations of hepatocytes, two populations of HSCs, two populations of macrophages, a population of cholangiocytes, and a population of LSECs. The gene expression profiles of the snRNA-seq clusters identified with this analysis had expression profiles consistent with the corresponding MERFISH clusters, supporting their assignment (**Supplementary Fig. 4b-d**). In addition, the snRNA-seq data identified a small fourth population of hepatocytes, a population of monocytes, and a population of lymphocytes, which we did not identify in the MERFISH data.

**Figure 3.**
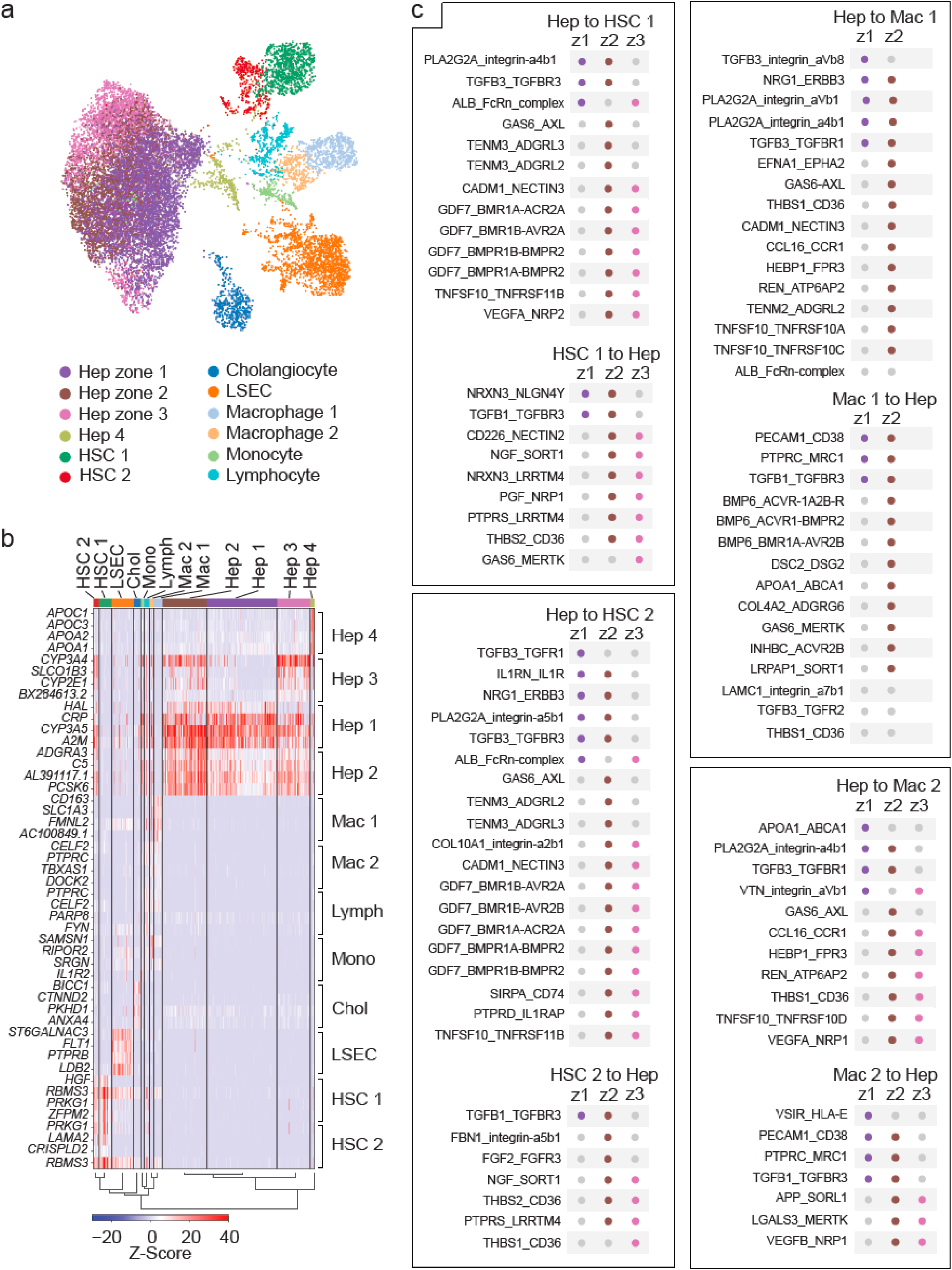
Combining MERFISH and snRNA-seq to define spatial interactions. **a.** UMAP of snRNA-seq data for healthy human liver. Annotations for the hepatocytes were determined by merging data from both MERFISH and snRNA-seq and leveraging both marker gene expression and label-transfer of MERFISH labels. **b.** Heatmap displaying differentially expressed genes that identify each cluster of snRNA-seq nuclei. Relative gene expression is indicated by z-score. Each column represents a single cell, and cells are grouped by cluster as indicated at the top of the heatmap. Genes are indicated to the left of the heatmap, and groups of genes characteristics of individual cell clusters are indicated on the right. Hierarchical clustering is shown at the bottom. **c.** Putative receptor-ligand interactions identified between zonal hepatocytes and notable non-parenchymal cells. Receptor-ligand interactions were mapped between each zone of hepatocytes (zone (z) 1, z2, z3) and cell types found in proximity to these cells based on MERFISH. Only interactions that are uniquely enriched by hepatocyte zonation are shown. Gray circles indicate no interaction, and colored circles (zone 1 hepatocytes - purple; zone 2 hepatocytes - brown; zone 3 hepatocytes - pink) indicate receptor ligand co-expression.

By investigating both the expressed genes and the cluster labels provided by MERFISH, we were able to associate three of the joint hepatocyte clusters with zones 1, 2, and 3 in the MERFISH data, and we assigned these joint cluster numbers to match their association with the classic zones (**Fig. 3a; Supplementary Fig. 4b, c**). The fourth hepatocyte cluster, not seen with MERFISH, contained 290 cells (2.7% of hepatocytes) and was defined by expression of *APOC1*, *APOC3*, *APOA1*, and *APOA2 (***Fig. 3b; Supplementary Fig. 4b**), suggesting increased activity in cholesterol and triglyceride metabolism^29^. These genes were not included in our MERFISH library, which may be why this population was not identified with MERFISH.

### Cell-cell interactions in space

While we did not observe clear zonal distributions in non-parenchymal cells to match hepatocytes, we noted that the variation in gene expression in hepatocytes could nonetheless create a zonality to the interactions between hepatocytes and these other cell populations. To explore this possibility, we leveraged our jointly integrated MERFISH and snRNA-seq data to explore potential modes of cell-cell interactions that could be shaped by liver zonation. Specifically, we focused this analysis only on interactions that are uniquely enriched between hepatocytes of specific zones and the cell types in close proximity to these zonated hepatocytes (**Fig. 3c; Supplementary Fig. 4e, f**). In the periportal region, hepatocytes (zone 1 or zones 1 and 2) produced TGFB3, which can interact with TGFBR1 expressed in HSC 2, Mac 1, and Mac 2 cells. This signal may be further supported in HSC 2 cells by expression of the adapter TGFBR3, and TGFB3 can be recognized in latent form by Mac 1 cells expressing integrin avb8. PLA2G2A, a secretory phospholipase, can interact with integrins a4b1, a5b1, and aVb1 allowing periportal hepatocytes to signal to HSC 1, HSC 2, Mac 1, Mac 2, and LSECs. NGR1 expression in periportal hepatocytes can signal through ERBB3, expressed by both HSC 2 and Mac 1 cells, while IL1RN produced by periportal hepatocytes can antagonize IL1R expressed by HSC 2 cells. In contrast, HSCs and Macs produced more TGFB1 compared to hepatocytes, and expression of TGFBR3 enriched in periportal hepatocytes could help promote TGF-β signaling. Both Mac 1 and Mac 2 cells could also signal to periportal hepatocytes through expression of PCAM1 and PTRC, which bind CD38 and MRC1, respectively, while Mac 1 cells could signal to zone 2 hepatocytes through BMP6. Similarly, LSECs express PCAM1 and could signal to periportal hepatocytes through CD38.

In the pericentral region, hepatocytes could signal to HSC 1 and HSC 2 cells through expression of GDF7, which can be recognized by multiple BMP receptors in HSC 1 and HSC 2 cells, and expression of TNFSF10 by pericentral hepatocytes is recognized by TNFRSF11B, which is expressed by HSC 1 and HSC 2 cells. VEGFA, expressed by pericentral hepatocytes is recognized by NRP2, which is expressed by HSC 1 cells, while expression of PTPRD by pericentral hepatocytes is recognized by IL1RAP, expressed by HSC 2 cells. Mac 1 and Mac 2 cells can receive signals from CCL16, HEPB1, and TNFS10 expressed by pericentral hepatocytes, while signaling from CADM1 and TENM2 in pericentral hepatocytes is recognized by NECTIN3 and ADGRL2, respectively, which are expressed by Mac 1 cells. THBS1/2 are produced by HSCs and Mac 1 cells and can signal to CD36 expressed by pericentral hepatocytes. GAS6 produced by HSC 1 and endothelial cells can signal to pericentral hepatocytes through MERTK, while VEGFB produced by Mac 2 cells can be recognized by NRP1 expressed by pericentral hepatocytes. In contrast, Mac 1 cells produce INHBC and LAMC, which can be recognized by ACVR2B and integrin a7b1, respectively, expressed by pericentral hepatocytes. These findings highlight receptor-ligand interactions that may regulate cross-talk between hepatocytes and non-parenchymal cells in zone specific patterns.

### Effect of nuclear content on gene expression in hepatocytes

Hepatocytes have the capacity to replicate their nuclei without dividing, generating polyploid and multinucleated cells^12–14^. It is not clear if additional nuclear content leads to defined gene expression changes that could modulate function or to what degree zonation influences nuclear content in the human liver. As an image-based approach to transcriptomics, MERFISH naturally provides a direct measure of the nuclear content of each cell. To leverage this capability, we counted the number of nuclei observed in individual hepatocytes and correlated this property with aspects of gene expression and spatial location. Indeed, we observed both single and multinucleated hepatocytes in the MERFISH data (**Fig. 4a**) and found that approximately a third of hepatocytes were multinucleated (**Fig. 4b**). The distribution of single and multinucleated hepatocytes did not change across hepatocyte zones (**Fig. 4c, d**), and we did not observe any significant changes in expression of genes between hepatocytes with one or two nuclei as quantified by MERFISH (**Fig. 4e**). Nonetheless, the total RNA counts per cell increased with nuclei number (**Fig. 4f, left**) as did the cell area (**Fig. 4f, center**), such that multinucleated hepatocytes tended to be larger and contain proportionally more RNA transcripts such that the RNA density was not dependent on the number of nuclei (**Fig. 4f, right**). These results suggest that multinucleated hepatocytes may contain more RNA transcripts within proportionally larger cells, which may underscore an increased metabolic capacity. However, this increase does not appear to be associated with differential gene expression or location of hepatocytes within the lobule, suggesting that these cells do not have distinct functional roles in the healthy liver, at least within the pathways we explored with MERFISH.

**Figure 4.**
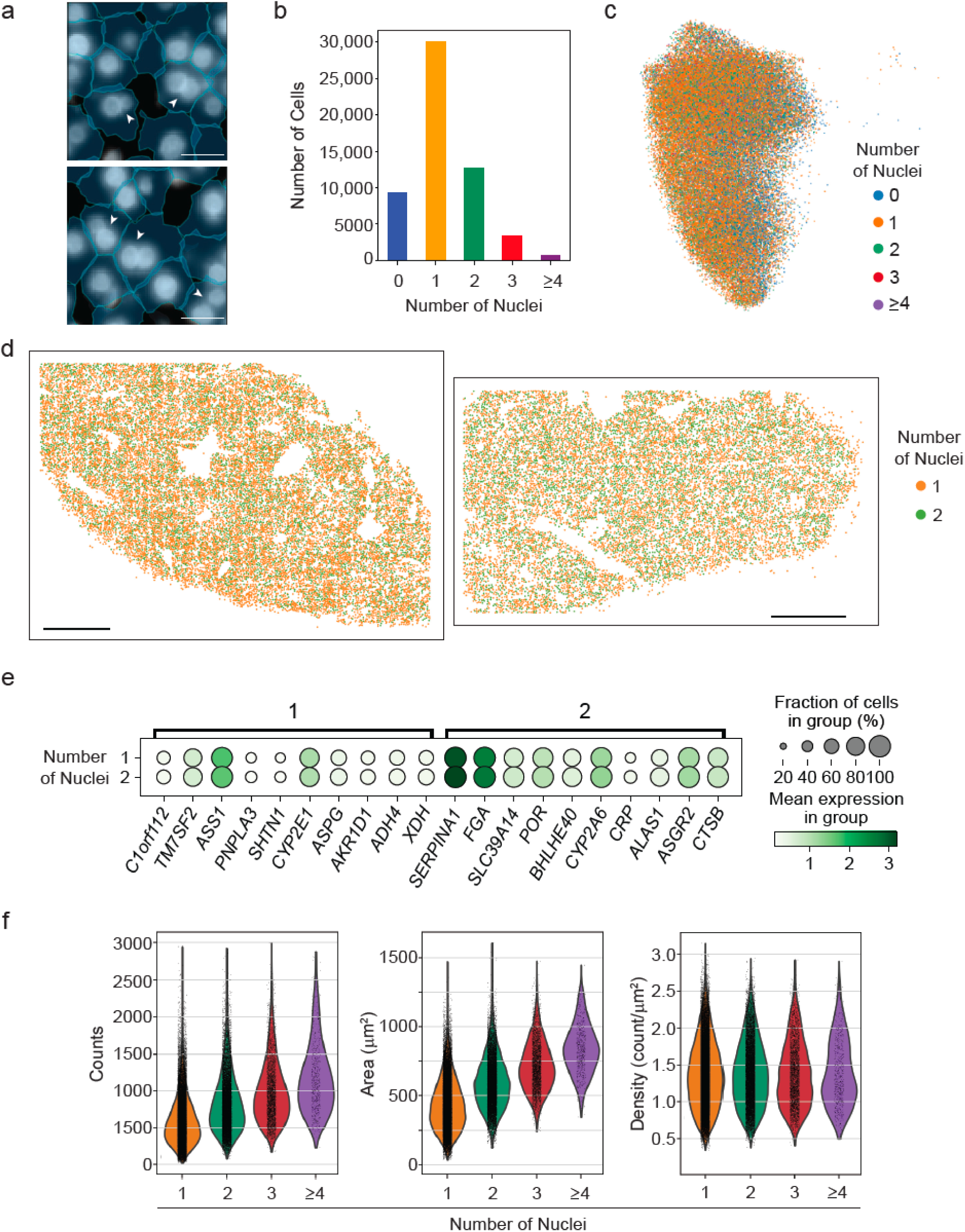
Multinucleated hepatocytes do not have unique spatial or gene expression distributions. **a.** Example MERFISH DAPI images showing hepatocytes containing single and multiple nuclei. Examples of multinucleated cells are indicated by white arrowheads. DAPI staining marks nuclei (white), and cell boundaries (light blue) were determined by Baysor with input from Cellpose. Scale bars: 20 μm. **b.** Distribution of number of nuclei per hepatocyte. **c.** Hepatocyte UMAP, as in Fig. 1, colored by the number of nuclei found within each hepatocyte. **d.** Spatial distribution of hepatocytes with 1 or 2 detected nuclei in healthy human liver (donor 1, left and donor 2, right). Scale bar: 1000 μm. **e.** Dot plots of hepatocyte genes for hepatocytes with 1 or 2 nuclei. The size of circles represents the percentage of cells expressing a gene in each group and the color intensity indicates mean expression. **f.** Distribution of the transcripts per cell (left), area per cell, (middle) and RNA density per cell (right) for hepatocytes with the indicated nuclear content.

### Applying spatial transcriptomics in liver fibrosis

In order to explore the spatial and cellular remodeling that occurs in the context of disease, we next performed MERFISH on human liver samples from patients with liver fibrosis (**Supplementary Table 1**). We leveraged the same protocols as described above to image and segment cells and, in total, imaged ∼460,000 cells. In order to compare hepatocytes between normal and fibrotic liver, we leveraged batch correction methods to jointly integrate MERFISH data across all samples **(Fig. 5a, b; Supplementary 5a, b**). The addition of hepatocytes from donors with fibrotic injury resulted in a periportal cluster of hepatocytes (Portal Hep) and a pericentral cluster of hepatocytes (Central Hep). Two new clusters were also identified that were almost entirely composed of hepatocytes from fibrotic injury (Fibrosis Hep 1 and 2). Periportal hepatocytes and pericentral hepatocytes retained their expected localization in the setting of chronic liver injury (**Fig. 5c**), while the two new clusters of hepatocytes that emerged with fibrotic injury were distributed throughout the lobules (**Fig. 5d**). A single cluster of HSCs and a single cluster of macrophages were also identified (**Fig. 5a, b**). HSCs displayed bands of cells across lobules, while macrophages were spread more diffusely through the lobules (**Fig. 5c**). These results identify expansion of two new populations of hepatocytes associated with development of fibrotic injury and a new bridging pattern of HSCs across the lobule. Thus we find in disease that hepatocyte zonation is at least partially maintained, and at two new injury-related hepatocyte states are greatly expanded that do not appear to retain zonation.

**Figure 5.**
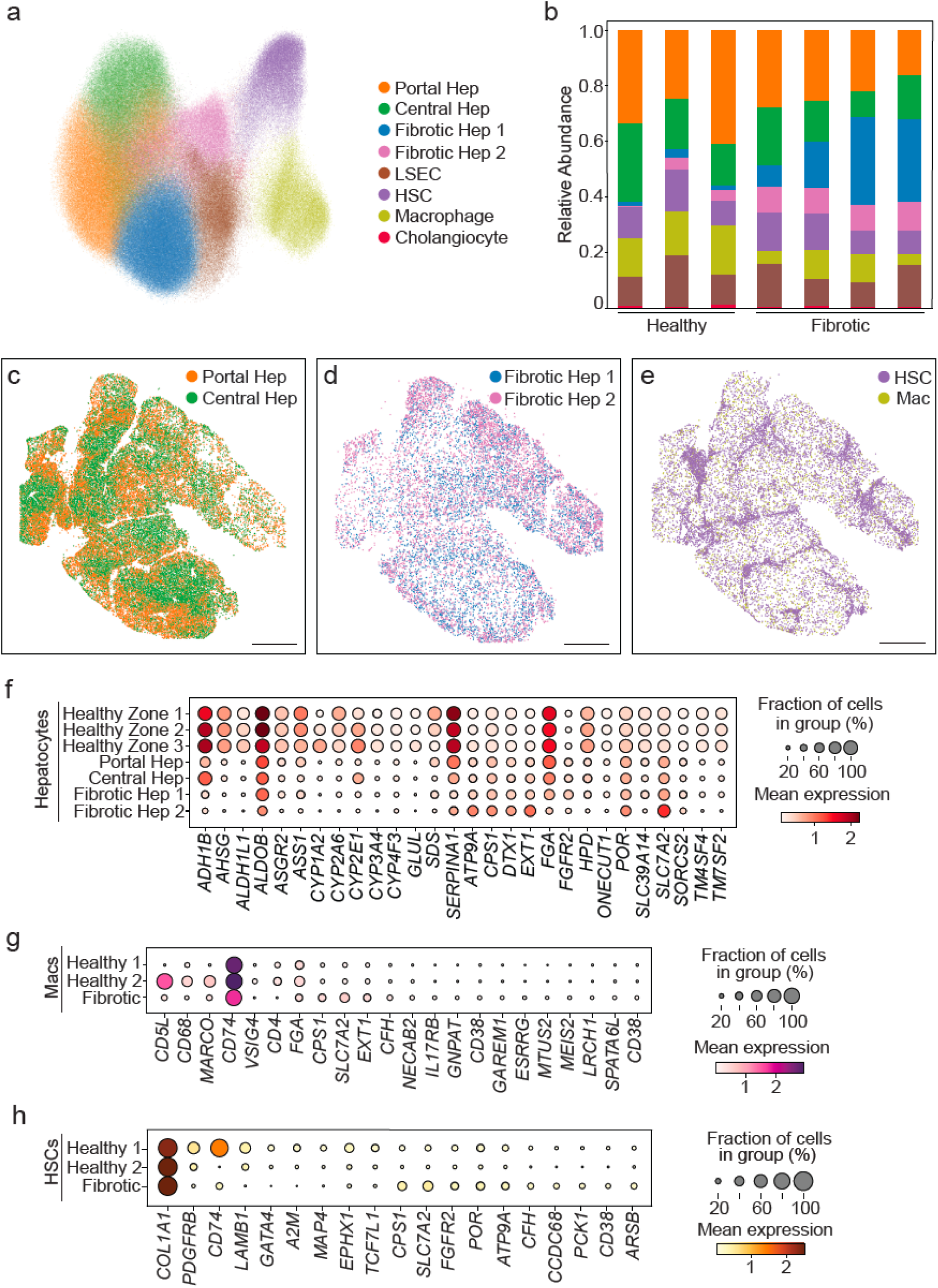
MERFISH reveals conserved zonation and new hepatocytes in fibrotic livers. **a.** UMAP of all cells measured with MERFISH including healthy samples and samples with fibrotic disease. Clusters are indicated by color. Portal Hep and Central Hep contain zone 1, 2, and 3 cells from healthy liver, and Fibrotic Hep 1 and Fibrotic Hep 2 contain new hepatocyte clusters not found in healthy liver. The HSC cluster contains both HSC 1 and HSC 2 cells from healthy liver, and the Mac cluster contains both Mac 1 and Mac 2 clusters from healthy liver. **b**. Relative abundance of each cell type identified in each sample. Cell type labels are the same as in **a**. **c**-**e.** Spatial distribution of cell populations in the fibrotic liver (donor 4), highlight hepatocytes that are similar to those seen in healthy samples (**c**), new hepatocyte populations that are identified only when including fibrotic liver samples (**d**), and key non-parenchymal cells (**e**). Scale bar: 1000 μm. **f-h**. Dot plot quantifying expression of genes differentially expressed in clusters seen in healthy measurements (Healthy zone 1, 2, 3) or in fibrotic samples (Portal Hep, Central Hep, Fibrotic Hep 1, Fibrotic Hep 2) for hepatocytes (**f**), macrophages (**g**), and HSCs (**h**). Macs Healthy 1 and Healthy 2 refer to Mac 1 and Mac 2 cells in healthy liver, and HSCs Healthy 1 and Healthy 2 refer to HSC 1 and HSC 2 cells in healthy liver.

We next evaluated changes in gene expression between healthy and fibrotic hepatocytes (**Fig. 5f**). For this analysis, we first separated healthy hepatocytes into their original three zones (Healthy zone 1, 2, 3) and compared expression to portal and central hepatocytes from fibrotic liver (Portal Hep and Central Hep). Portal and central hepatocytes from the fibrotic liver retain the same zonal expression of marker genes as healthy zone 1, 2, and 3 and are characterized primarily by a reduction in expression of these genes. We then evaluated gene expression of the newly expanded fibrotic populations (Fibrotic Hep 1 and 2) (**Fig. 5f, bottom**). The Fibrotic Hep 1 cluster showed a general reduction in expression of genes enriched in hepatocytes, while the Fibrotic Hep 2 cluster showed an increase in expression of several notable genes, including *SCL7A2*, *CPS1, ATP9A, DTX1, POR* and *EXT1* compared to all other hepatocyte populations. Fibrotic macrophages showed a loss of *MARCO*, *CD5L*, *CD68* expression characteristic of Mac 2 (noninflammatory) macrophages and retained an expression pattern that more closely matched Mac 1 cells in normal liver (**Fig. 5g**), while fibrotic HSCs maintained high levels of *COL1A1* expression without a clear positive signature in the available MERFISH probe sets (**Fig. 5h**). These results identify expansion of two hepatocyte populations with fibrotic injury that are not zonally distributed, suggesting that they may arise with equal frequency from all zonated healthy hepatocytes. Moreover, the increased expression of key genes, such as *CPS1* and *SCL7A2*, suggest that one of these hepatocyte states may be involved in increased urea cycle activity.

## Discussion

Significant advances in mapping the human liver at single-cell resolution have been achieved recently with scRNA-seq and snRNA-seq^8,9,25,30^. In parallel, platforms such as Visium^10^ now provide spatial information but not at single-cell resolution, making it challenging to assign transcripts to individual cells and cell types located in close proximity. Here we complement these previous efforts by applying MERFISH to achieve spatial transcriptomic analysis of >300 genes at single-cell resolution in the human liver, profiling nearly 600,000 cells in both healthy and diseased liver. Combining MERFISH with snRNA-seq then allowed us to analyze the broader transcriptome within spatially resolved cell types and cell sub-types.

From these measurements, we were able to resolve three clusters of hepatocytes that are spatially distinct across the lobule in healthy human liver, characteristic of separation of hepatocytes into zones 1, 2, and 3 moving from periportal to pericentral regions. While MERFISH allows to us to define these zones, it is also evident through pseudotime analysis, measurement of gene expression across individual lobules, and transcriptional analysis performed across many lobules, that zonation reflects a gradient from periportal to pericentral regions rather than distinct zones. Thus, we propose that the classic three zones are, in reality, an approximation for the continuous gene expression and functional changes that occur in space with human hepatocytes--a proposal consistent with recent single-cell analysis in the murine liver^23,24^.

With the development of fibrosis and cirrhosis, many hepatocytes maintained zonal phenotype, but gene expression was reduced as quantified by MERFISH. This could represent a reduction in gene expression with injury or reflect a technical challenge related to adapting the protocol to fibrotic tissue. We also observed the emergence of two additional hepatocyte populations with cirrhosis (Fibrotic Hep 1 and 2 populations; **Fig. 5a, b, d**). Fibrotic Hep 1 cells showed diffuse distribution through the lobules without a portal or central bias and were primarily characterized by suppressed expression of zonal genes. Fibrotic Hep 2 cells also did not exhibit zonal distribution and showed notable induction of *CPS1* and *SCL7A2*. CPS1 (Carbamoyl phosphate synthetase I) is responsible for converting ammonia to carbamoyl phosphate to enter the urea cycle^31^ while *SCL7A2* encodes an arginine/ornithine/lysine transporter and provides ornithine (either directly) or through conversion of arginine to interact with carbamoyl phosphate to form citrulline^32,33^. The increase in expression of these genes in Fibrotic Hep 2 cells may reflect the ability of these cells to process increased ammonia levels that often develop in end-stage liver disease as a result of reduced activity in hepatocytes more globally^34,35^ or serve to activate bone marrow-derived monocytes through increased *CPS1* expression^36^.

Two populations of macrophages were identified by MERFISH and show different distribution patterns through the lobule. Mac 2 cells were identified as *MARCO*^+^*, CD5L*^+^, and *CD68*^+^ with increased expression of *VISG4* compared to Mac 1 cells and were located more diffusely through the lobule. Mac 1 cells were *MARCO*^-^ and *CD5L*^-^ with lower levels of *CD68*. While Mac 1 cells were also found through the lobule, they also exhibited a distinct enrichment in portal areas in contrast to Mac 2 cells. With the development of fibrosis, the macrophage population lost expression of *MARCO*, *CD5L*, and *CD68* suggesting a shift to a Mac 1 phenotype with fibrotic injury based on the available markers from MERFISH.

MERFISH also identified two populations of HSCs. HSC 1 was enriched in the portal areas compared to HSC 2, and both populations were found distributed through the lobules. Within the genes evaluated by the MERFISH probes, the MHC class II chaperone *CD74*^37^ showed the greatest differential expression between the two HSC populations, with highest expression observed in HSC 1 cells. *CD74* is expressed by HSCs, can be induced by inflammatory signals such as IFN-γ and can promote immune activation through MHC class II-mediated interactions^38,39^. It is currently unclear if these *CD74* positive and negative populations represent HSCs with distinct antigen presentation abilities or possibly different histories of activation.

With single-cell spatial resolution, we evaluated receptor-ligand coexpression in cells across hepatocyte zonation. Our MERFISH panel was not designed to explore a diverse set of functional pathways in HSCs, macrophages, or LSECs across zones, but the current data do allow us to ask about specific interactions driven by differential expression of receptors or ligands in hepatocytes across zones. Indeed, we clearly see that the zonal expression of receptors and ligands in hepatocytes can reshape the potential interactions with cells in which we see no obvious zonal expression differences in our data. In particular, we identified signaling interactions including a thrombospondin signal (THSB1/2) from HSCs and macrophages to CD36 in zone 3 hepatocytes, which may influence metabolic dysfunction associated steatotic liver disease (MASLD)^40^, and an interaction between PECAM1, expressed by macrophages and LSECs, and CD38 enriched in zone 1 hepatocytes, which could affect gluconeogenesis^41^.

MERFISH also provides data about nuclei and cell boundaries in addition to gene expression, allowing us to link nuclei content to gene expression within individual cells. We found that approximately a third of adult hepatocytes are multinucleated, in agreement with previous descriptions of human liver. We did not quantify ploidy separately, but based on these studies, we would predict that >80% of these multinucleated hepatocytes are 2N^42,12^. While we did not observe a change in relative gene expression with increase in nuclear content, we did observe an increase in RNA transcripts. In addition, multinucleated hepatocytes also tend to be larger in size such that mRNA levels were proportional to cell size. These findings are consistent with the interpretation that cells with more nuclei are both larger^13^ and express higher levels of RNA. Most of the transcripts identified in hepatocytes are cytoplasmic, but transcripts are also present in the nucleus. It is possible that MERFISH data could be used to evaluate transcriptional activity between different nuclei in the same hepatocyte, but this will require additional tools to reconstruct nuclei from the stacked images and control for the fraction of each nuclei visualized in an individual cell.

Taken together, our study applied MERFISH to create spatial transcriptomic maps of the healthy and fibrotic liver at single-cell resolution to define spatially and transcriptionally distinct sub-populations of hepatocytes, macrophages, and HSCs within hepatic lobules and distinct hepatocyte sub-populations that expand with fibrotic injury. By combining MERFISH with snRNA-seq, we extended the transcriptional profiles of cell types in these spatial maps to understand unique receptor-ligand interactions involving hepatocytes and non-parenchymal cells across hepatocyte zones. Finally, by evaluating nuclear content, we found that multinucleated hepatocytes do not show differences in relative gene expression compared to mono-nucleated hepatocytes, but tend to be larger and produce more total transcripts. Future studies will now be able to extend these approaches, expanding probe set libraries to encompass additional RNA species to evaluate all cell types in the liver and their broader transcriptional profiles at single-cell spatial resolution.

## Acknowledgments

The authors thank Eliana Epstein and Jasneet Aneja for organizing the collection of liver tissue and Nicole Brousaides and the MGH Histopathology Research Core for help sectioning tissue. We thank Sonya MacParland, Gary Bader, Tallulah Andrews, and Jawairia Atif for helpful discussions and assistance in selecting genes for the MERFISH probe set. We thank Cristin McCabe for assistance with data management, and Adam Slamin, Dan Dubinsky, and the Broad Genomics Platform for help with generation of single-nucleus sequencing data. R.J.X., J.R.M., and A.C.M were supported by grants from the Chan Zuckerberg Initiative.

## Author contribution

B.P., B.W., J.R.M, and A.C.M. conceived the study and designed the experiments. Samples were prepared for MERFISH by B.P. and imaged by B.W. with support from A.S. snRNA-seq was performed by L.A-Z with support from A. S., J.D., and R.J.X. Computational analyses were performed by B.W. and B.P. with support from R.U.R. The manuscript was written by B.W., B.P., J.R.M, and A.C.M. with input from all other authors.

## Competing Interest/conflict of interest

R.J.X. is a co-founder of Celsius Therapeutics and Jnana Therapeutics, board director at MoonLake Immunotherapeutics, and consultant to Nestlé. J.R.M is a co-founder of, stake-holder in, and advisor for Vizgen. J.R.M. is an inventor on patents associated with MERFISH applied for on his behalf by Harvard University and Boston Children’s Hospital. J.R.M.’s interests were reviewed and are managed by Boston Children’s Hospital in accordance with their conflict-of-interest policies. A.C.M. receives research funding from Boehringer Ingelheim and GlaxoSmithKline for unrelated projects.

## Methods

### Tissue collection

All liver samples were collected from excess surgical tissue in accordance with protocols approved by the Mass General Brigham Institutional Review Board (IRB).

### Sample preparation

Liver tissue sections (3-5 mm thickness) were fixed by submerging in fresh 4% v/v paraformaldehyde (PFA; EMS 15710) in 1X Phosphate Buffered Saline (PBS; Ambion AM9625) for 3 - 5 hours at 4 °C with gentle rocking. After fixation, tissue samples were then transferred to a 30% w/v sucrose (VWR, 0335) solution in 1X PBS supplemented with 4% v/v PFA and incubated overnight at 4 °C with gentle rocking. After washing in 1X PBS, samples were placed in cryomolds on dry ice, covered with Optimal Cutting Temperature media (OCT; Tissue-Tek 4583), wrapped in aluminum foil, and stored at -80 °C until use.

### MERFISH library preparation

A panel of 317 genes were selected with a focus on those differentially expressed between hepatocyte clusters^10^ and also included common markers of non-parenchymal liver cells^8^. The MERFISH encoding probes targeting these genes were designed using a previous pipeline^17,43^. Briefly, each probe contained a 30-nt-long region specific to the targeted RNA, concatenated with a series of readout sequences that defined the binary barcode assigned to that RNA. Each mRNA was targeted with 72 such probes. To encode these 317 genes, we selected a 22-bit-long, constant Hamming weight and Hamming distance code with a Hamming weight of 4 and a minimum Hamming distance of 4. Each of the 72 encoding probes contained two readout sequences and the complement of all encoding probes to a given gene contained instances of the four readouts associated with the 4 bits in which they had a ‘1’. This barcoding scheme contains 332 barcodes and we leveraged the 15 not used to encode RNAs as ‘blank’ barcode controls, not assigned to an RNA.

Templates for these probes were designed by concatenating a primer and a T7 promoter to the sequences, as described previously, and the template pool was ordered from Twist Biosciences. These templates were amplified into encoding probes in a protocol that involved PCR, *in vitro* transcription, reverse transcription, alkaline hydrolysis, and SPRI purification, as described previously^17,18,43,44^.

### MERFISH staining

The general workflow for MERFISH sample preparation includes sectioning, permeabilization, cell boundary staining, RNA probe hybridization, acrylamide gel embedding, digestion, and photobleaching of samples. These protocols have been described previously^17,18,43,44^. Briefly, OCT-embedded tissues were cryosectioned at a thickness of 7 μm, and slices were transferred onto silanized, poly-lysine coated 40-mm circular coverslips with fiducial beads, prepared as described previously^17,18,43,44^. Following sectioning, tissue slices were briefly fixed with 4% v/v PFA in 1X PBS at room temperature for 10 minutes, washed thrice with 1X PBS for 5 minutes, and then permeabilized overnight at 4 °C in 70% ethanol. Samples were rehydrated in 2X saline sodium citrate (SSC; Ambion AM9763).

To prepare samples for antibody staining of cell boundaries, tissue samples were treated with 0.05% (v/v) proteinase K (New England Biolabs [NEB]; P8107S) in pre-warmed (37 °C) 2X SSC for 10 minutes at 37 °C. Samples were then rinsed with 2X SSC before incubating with a blocking buffer (10% BSA, 3% v/v 6% v/v murine RNase inhibitor [NEB, M0314L] in 2X SSC) for 30 minutes. The sample was then stained with a primary antibody for Na/ATPase (2 µg/ml, Abcam; 76020) in the same blocking buffer for 30 minutes at room temperature. Excess primary antibody was removed from the samples with three, 10-minute, 2X SSC washes at room temperature. The sample was stained with a secondary antibody using the same protocol as the primary antibody (3.75 µg/ml; ThermoFisher; A16112). The secondary antibody was labeled with an oligonucleotide using previous protocols^45^.

Probes were hybridized to the sample at a concentration of 6-10 µM in a 30% v/v formamide (Fisher Scientific, AM9342), 10% w/v dextran sulfate (VWR, 97062-828), 1 mg/mL yeast tRNA (ThermoFisher, 15401029) solution in 2X SSC for 48 hours at 37 °C in a humidified oven. Samples were rinsed by letting the samples sit in 30% v/v formamide in 2X SSC before hybridization for at least 4 hours and twice after hybridization for 30 minutes each time at 37 °C.

To clear samples, they were embedded in a polyacrylamide gel and then digested for two days with proteinase K, as described previously^17,18,43^. Briefly, samples were embedded in a thin polyacrylamide film by inverting them onto a GelSlick-coated microscope slide with a droplet of 4% acrylamide solution (4% v/v 20:1 acrylamide:bis-acrylamide [BioRad, 1610144] with 0.15% v/v TEMED [Sigma, T7024] and 0.30% v/v ammonium persulfate [Sigma, 215589]). After the gel polymerized for 2 hours, samples were covered with a digestion solution (1% proteinase K, 20% v/v SDS [ThermoFisher, AM9823], 0.25% v/v triton-X [Sigma, T8787] in 2X SSC) and incubated at 37 °C for 24 hours. Samples were then rinsed multiple times with 2X SSC. In order to remove autofluorescence, samples were photobleached under a blue LED light at 4 °C for 24 hours. Samples were stored at 4 °C until imaging.

### snRNA-seq

Single nucleus extraction was performed based on previously-described protocols^46^. A 2X stock of salt-Tris solution (ST buffer) containing 292 mM NaCl (Thermo Fisher Scientific, AM9759), 20 mM Tris-HCl pH 7.5 (Thermo Fisher Scientific, 15567027), 2 mM CaCl_2_ (VWR International Ltd, 97062-820), and 42 mM MgCl_2_ (Sigma Aldrich, M1028) was prepared in ultrapure water. The day of each experiment, for each sample, Tween with salts and Tris (TST) buffer was made from 1 ml of 2X ST buffer, 60 µl of 1% Tween-20 (Sigma Aldrich, P-7949, 0.03% final), 10 µl of 2% BSA (NEB, B9000S, 0.01% final) and 930 µl of nuclease-free water and supplemented with 1U/ml Protector RNase inhibitor (Millipore Sigma, 3335402001). 1X ST buffer was prepared by dilution 2X ST with ultrapure water (Thermo Fisher Scientific, 10977023) and supplemented with 0.5U/ml Protector RNase Inhibitor (Millipore Sigma, 3335402001).

On dry ice, a section of frozen tissue was placed into a gentleMACS C Tube (Miltenyi Biotec, 130-093-237) with 2 ml of TST buffer. Tissue was immediately dissociated using a gentleMACS Dissociator (Miltenyi Biotec, 130-096-427) using the m_Spleen_01_01 program twice and incubated on ice for 5 min to complete a 10 min incubation in Tween with salts and Tris (TST) buffer. C tubes were centrifuged at 4 °C for 2 min at 500*g*. The pellet was resuspended in TST buffer, filtered through a 40 µm Falcon cell strainer (VWR, 43-50040-51) into a 50 ml conical tube. The strainer was washed with 1 ml 1X ST buffer before use. An additional 1 ml of 1X ST buffer was used to wash the gentleMACS C Tubes and filter, and another 1 ml was added to the filter for a final wash. The sample was transferred to a 15 ml conical tube and centrifuged at 4 °C for 10 min at 500*g*. The pellet was resuspended in 1X PBS (-Mg/-Ca, Gibco, 10010023), 1% BSA (NEB, B9000S), and 1U/ml Protector RNase Inhibitor (between 100-200 μl depending on pellet volume). The nucleus solution was filtered through a 35 µm Falcon cell strainer (Corning, 352235). Nuclei were counted using a INCYTO C-chip disposable hemocytometer (VWR, 22-600-100).

### Sample processing

#### MERFISH measurements

MERFISH measurements were performed as described previously using a custom-microscope system^44^. The entire slice was first imaged using a 10X objective and the DAPI stain, and this mosaic was used to select the region of interest for MERFISH measurements.

#### snRNA-seq

8,000 - 12,000 nuclei of the single nucleus suspension were loaded onto the Chromium Chips for the Chromium Single Cell 3′ Library (Chromium Next GEM Single Cell 3’ Kit v3.1; PN-1000268, PN-1000120, PN-1000215) according to the manufacturer’s recommendations (10x Genomics). Gene expression libraries were constructed and indexed according to manufacturer’s instructions and pooled for sequencing on a NovaSeq 6000 sequencer (Illumina). All libraries were sequenced to a targeted depth of 400 million reads in the following configuration: R1: 28 bp; R2: 90 bp; I1, I2: 10 bp.

### Analysis

#### MERFISH decoding

MERFISH decoding was performed using a previously described pipeline^17,18,43^. Briefly, images from a single field-of-view (FOV) were stacked into movement-corrected stacks containing all z-planes and imaging channels, an optimal weighting was determined for the relative intensity of different image frames, and pixels were assigned to individual barcodes using a soft decoding approach based on nearest neighbors with a Euclidean distance metric. Adjacent pixels assigned to the same barcode were aggregated to form a single RNA.

RNAs were assigned to cells using a combination of Cellpose^20^ (version 0.7.2) and Baysor^19^(version 0.5.0). Briefly, we trained a cellpose model based on the Na/K ATPase immunofluorescent stain, and created a label matrix using cellpose and this model for each FOV. Cellpose was applied to each z-plane separately, and then overlapping boundaries were combined across z-planes to create a 3D segmentation. We leveraged our previous pipeline^17,18,43^ to identify cell boundaries, connect boundaries across FOV boundaries, and parse RNAs into these boundaries. This initial segmentation result was then refined and improved with Baysor using the following parameters: scale 10, scale standard deviation 50%, molecules/cell 3, and prior confidence 0.50.

#### MERFISH single-cell analysis

The output of Baysor was analyzed using Scanpy^47^ (version 1.8.1). Initial processing concatenated the MERFISH data into a single AnnData structure, filtered cells with less than 3 genes and 15 RNA counts, normalized the remaining cells, and applied a logarithmic scaling. The processed cells were analyzed by PCA with 100 components. Harmony was used as our data integration method in order to account for sample variation^48^. Cells were clustered with a variety of leiden resolutions; the final resolution for downstream analysis was set at 1.1.

#### snRNAseq

Samples were demultiplexed using Illumina’s bcl2fastq conversion tool and the 10x Genomics pipeline Cell Ranger (version 6.0.1) to perform alignment against the 10x Genomics pre-built Cell Ranger reference GRCh38-2020-A (introns included), filtering, barcode counting, and UMI counting. The SoupX package^49^ (version 1.6.2) was used to remove ambient RNA contamination for each individual sample. Count data was then filtered, clustered, and represented in low-dimensional embedding UMAP using Scanpy^47^ (version 1.9.1). ). Cells were retained if they expressed at least 500 genes and contained <5% mitochondrial reads. Genes were retained if they were present in at least 10 cells. After applying these filtering steps on 16,662 input nuclei, the dataset contained 15,278 high-quality single nuclei that were eligible for further analysis. Receptor-ligand interactions were evaluated with CellPhoneDB^50,51^. We applied 1000 iterations and considered ligands and receptors for analysis if expressed by at least 10% of cells (threshold = 0.1). Predicted receptor-ligand interactions were excluded if 1) the ligand was expressed at a higher level in the receptor cell than the ligand cell, 2) the ligand was not a single gene product, 3) the interaction score for a significant interaction was <2% greater than the interaction score for non-significant interaction between hepatocytes and the indicated cell type.

#### MERFISH-snRNAseq Integration

To co-integrate the healthy MERFISH and snRNA-seq measurements, we concatenated the two separate AnnData structures after each dataset was normalized by cell counts. Expression was logarithmically transformed, z-scored, and the genes were trimmed to just those measured with MERFISH. These cells were then integrated using Harmony in order to compare gene expression markers and labels across cell types, leveraging the inbuilt algorithms and defaults associated with ScanPy. Co-integrated data were clustered using Leiden clustering. The individual clusters were then labeled by leveraging a combination of marker expression and the existing annotations for the MERFISH cells derived from the clustering analysis of the MERFISH healthy data alone as described above. Importantly, this approach was used only to apply labels to the snRNAseq cells, and all expression analysis derived from this co-integration was taken from the snRNAseq cells.

