## Supplementary Figures for "Spatial transcriptomics of healthy and fibrotic human liver at single-cell resolution"

| Donor | Analysis ID | Age | Gender | Condition | snRNA-seq Designation |
|---|---|---|---|---|---|
| 1 | AM042 | 51-60 | F | Healthy | 6854-13 |
| 2 | AM061 | 81-90 | F | Healthy | 6854-18 |
| 3 | AM048 | 61-70 | M | Healthy | 6854-14 |
| 4 | AM031 | 51-60 | M | Fibrosis (F4) | |
| 5 | AM062 | 41-50 | F | Fibrosis (F2) | |
| 6 | AM066 | 31-40 | F | Fibrosis (F4) | |
| 7 | AM072 | 61-70 | M | Fibrosis (F4) | |

**Supplementary Table 1. Summary of human liver samples.**

Sample is a unique number assigned to each sample. Analysis ID represents a unique identifier assigned to samples during collection. Age is the age range of the donor. Gender is the gender of the donor. Condition indicates the diagnosis of the liver sample. F2 indicates metavir fibrosis score of 2; F4 indicates a metavir fibrosis score of 4 (cirrhosis). snRNA-seq Designation indicates the ID assigned to each sample in the snRNA-seq data deposited on GEO.

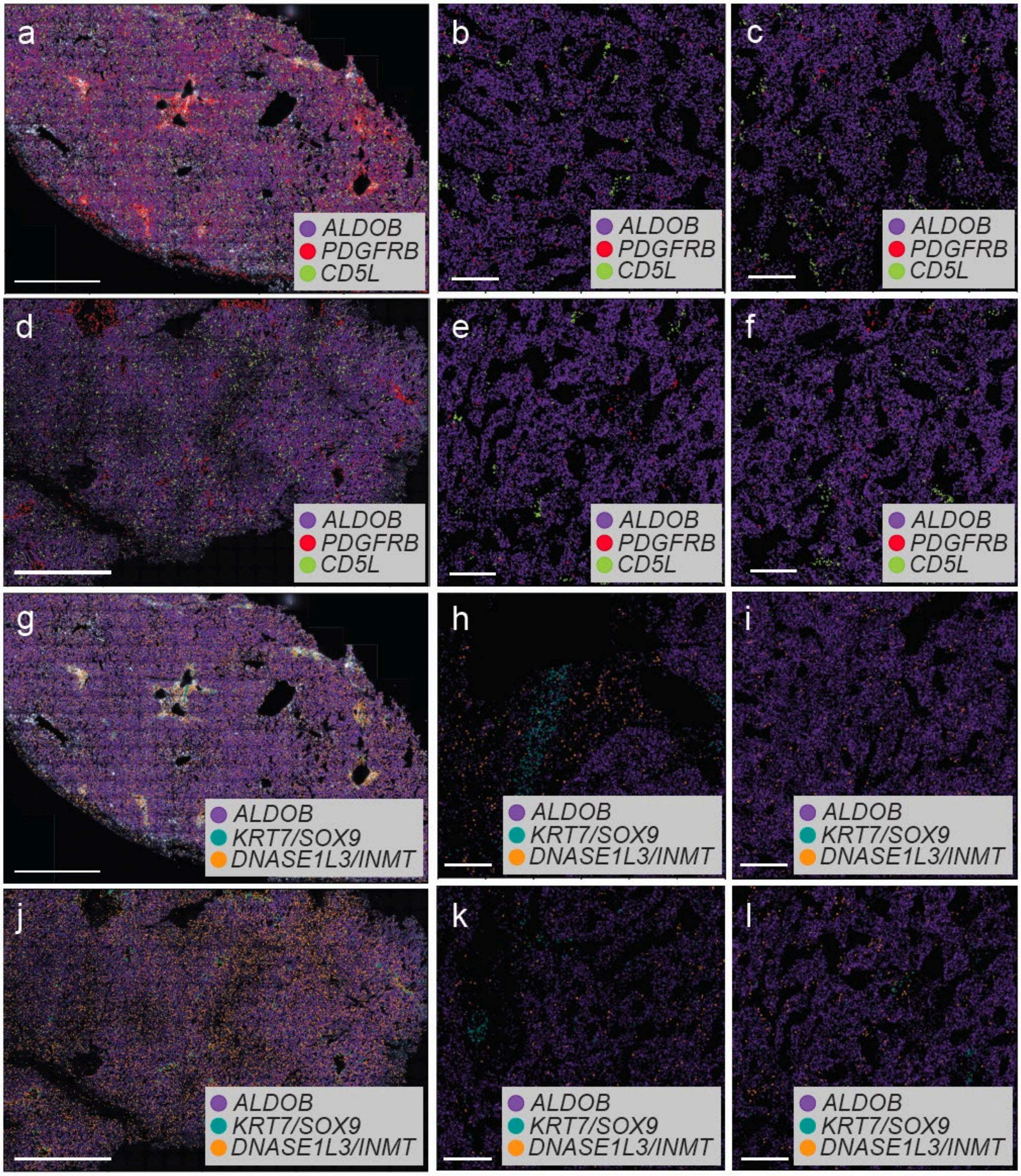
a
ALDOB
PDGFRB
CD5L
b
ALDOB
PDGFRB
CD5L
c
ALDOB
PDGFRB
CD5L
d
ALDOB
PDGFRB
CD5L
e
ALDOB
PDGFRB
CD5L
f
ALDOB
PDGFRB
CD5L
g
ALDOB
KRT7/SOX9
DNASE1L3/INMT
h
ALDOB
KRT7/SOX9
DNASE1L3/INMT
i
ALDOB
KRT7/SOX9
DNASE1L3/INMT
j
ALDOB
KRT7/SOX9
DNASE1L3/INMT
k
ALDOB
KRT7/SOX9
DNASE1L3/INMT
l
ALDOB
KRT7/SOX9
DNASE1L3/INMT

**Supplementary Figure 1. Visualizing RNA transcripts that define individual cell types with MERFISH. a** The spatial distribution of mRNAs for *ALDOB* (marker of hepatocytes), *PDGFRB* (marker of HSCs), and *CD5L* (marker of macrophages/Kupffer cells) in tissue from donor 1. Scale bar: 1000 µm. **b,c.** Magnified image from tissue section shown in (**a**)**.** *ALDOB* (purple) maps to hepatocytes, and *CD5L* (green) maps to macrophages. *CD5L* transcripts cluster in the sinusoids (black space between hepatocytes) consistent with sinusoidal localization. *PDGFRB* (red) maps to HSCs, and transcripts are found along the edge of hepatocytes, consistent with subendothelial localization. Scale bar: 50 µm. **d-f.** Spatial distribution as in (**a-c**) repeated in liver tissue from donor 2. **g.** The spatial distribution of mRNAs for *ALDOB* (hepatocytes), *KRT7/SOX9* (cholangiocytes), and *DNAS1L3/INMT* (LSECs). **h, i.** *KRT7/SOX9* (green) identifies cholangiocytes clustered in portal regions (**h**), and *DNAS1L3/INMT* (red) identifies LSECs spread diffusely (**i**) along the edges of hepatocytes (purple). **j-l**. Analysis shown in (**g-i)** repeated in tissue from donor 2.

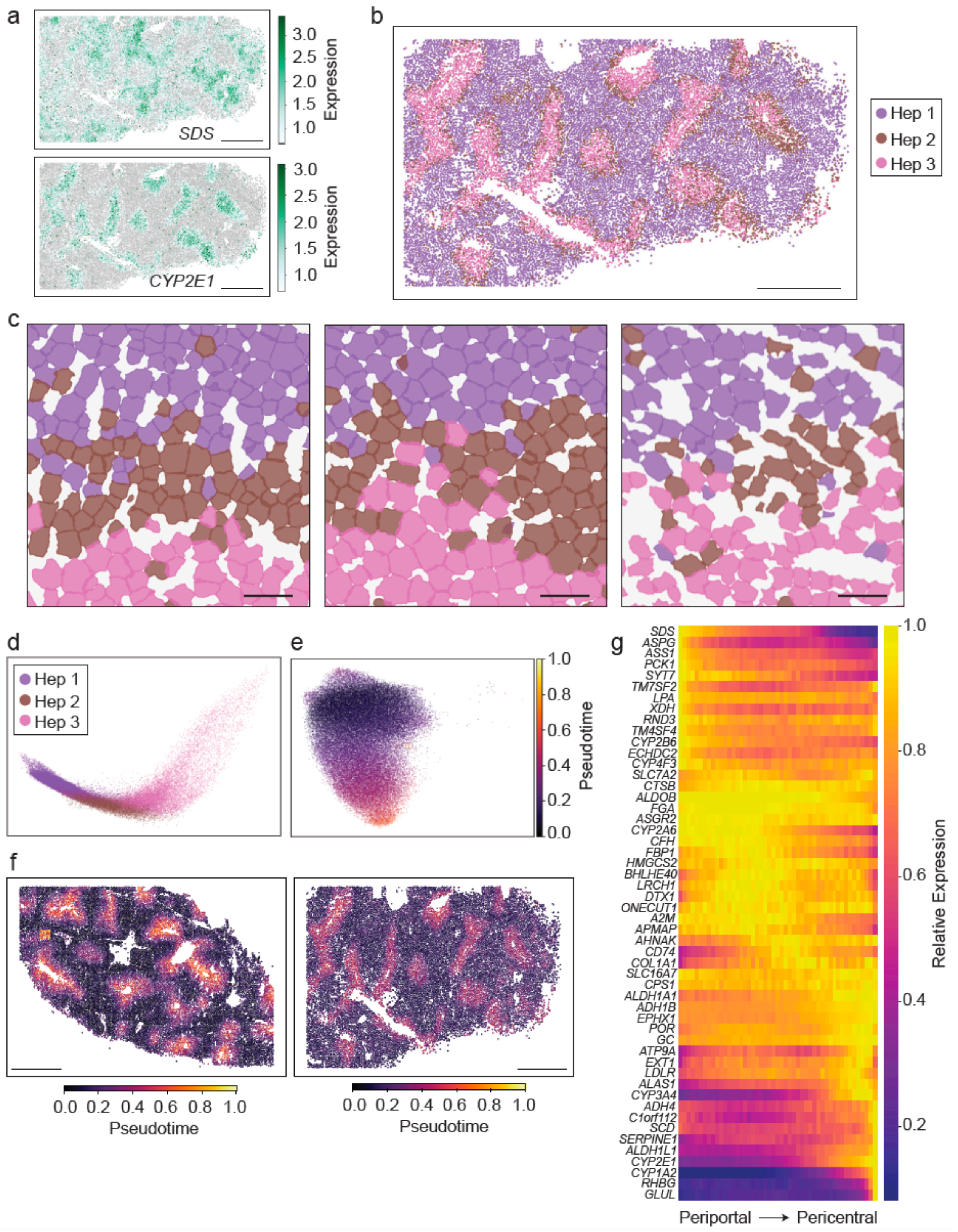
a
SDS
CYP2E1
Expression
3.0
2.5
2.0
1.5
1.0
b
Hep 1
Hep 2
Hep 3
c
d
e
Pseudotime
f
g
Relative Expression
Periportal ⟶ Pericentral
SDS
ASPG
ASS1
PCK1
SYT7
TM7SF2
LPA
XDH
RND3
TM4SF4
CYP2B6
ECHDC2
CYP4F3
SLC7A2
CTSB
ALDOB
FGA
ASGR2
CYP2A6
CFH
FBP1
HMGCS2
BHLHE40
LRCH1
DTX1
ONECUT1
A2M
APMAP
AHNAK
CD74
COL1A1
SLC16A7
CPS1
ALDH1A1
ADH1B
EPHX1
POR
GC
ATP9A
EXT1
LDLR
ALAS1
CYP3A4
ADH4
C1orf112
SCD
SERPINE1
ALDH1L1
CYP2E1
CYP1A2
RHBG
GLUL

**Supplementary Figure 2. Hepatocyte zonation in healthy human liver is a continuous process. a.** Spatial distribution of cells from donor 2 colored by expression of *SDS* (enriched in zone 1) and *CYP2E1* (enriched in zone 3). Normalized expression level is indicated (far right). Scale bars: 1000 μm. **b.** Cells assigned to each cluster of hepatocytes (Hep 1, Hep 2, Hep 3) are shown within the architecture of a liver section from a second donor. Hep 1 cells (purple) map to periportal areas (zone 1), Hep 3 cells map to pericentral areas (zone 3), and Hep 2 cells map between Hep 1 and Hep 3 cells (zone 2). Scale bar: 1000 μm. **c.** Spatial distribution of hepatocytes from three individual donors (donor 1, left, donor 2, center, donor 3, right). Zone 1 hepatocytes are shown in purple, zone 2 hepatocytes in brown, and zone 3 hepatocytes in pink. Cell boundaries are depicted as polygons. Scale bars: 50 μm. **d.** Diffusion coefficient analysis of healthy hepatocyte clusters colored by cluster label. **e.** Pseudotime of hepatocytes projected onto the UMAP plot of hepatocytes. **f.** Pseudotime projected into tissue sections from donor 1 (left) and donor 2 (right) and showing transition from zone 1 to zone 3 with increasing pseudotime. Scale bars: 1000 μm. **g.** Average gene expression within hepatocytes versus hepatocyte pseudotime.

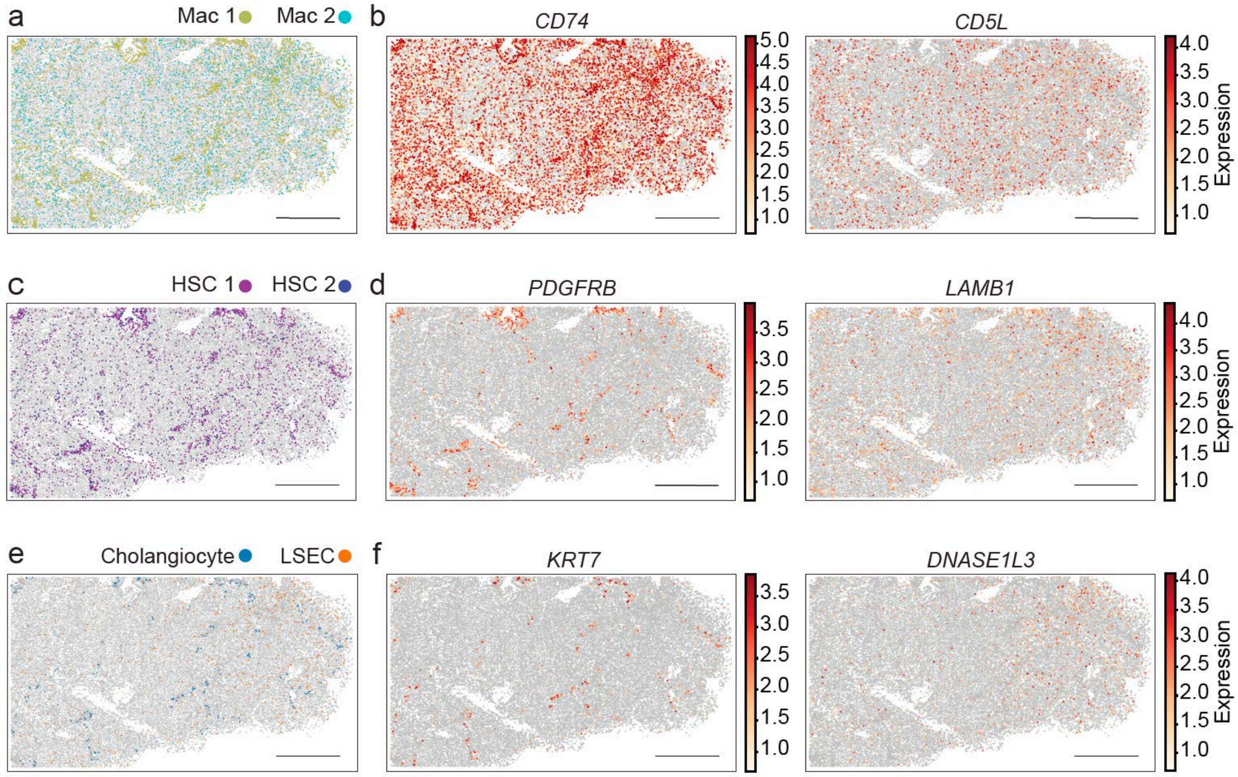


**Supplementary Figure 3. MERFISH reveals the distribution of non-parenchymal cells in healthy human liver. a,c,e**. Spatial distribution of macrophage (**a**), HSC (**c**), and cholangiocyte and LSEC (**e**) populations mapped in the same tissue section as **Fig. 2b** (donor 2**).** Scale bars: 1000 µm. **b, d, f**. Spatial distribution of cells in tissue section from donor 2 (**Fig. 2b**) colored by the normalized expression of the indicated marker genes. *CD74* and *CD5L* identify macrophages, *PDGFRB* and *LAMB1* identify HSCs, *KRT7* identifies cholangiocytes, and *DNAS1L3* identifies LSECs. Scale bars: 1000 µm.

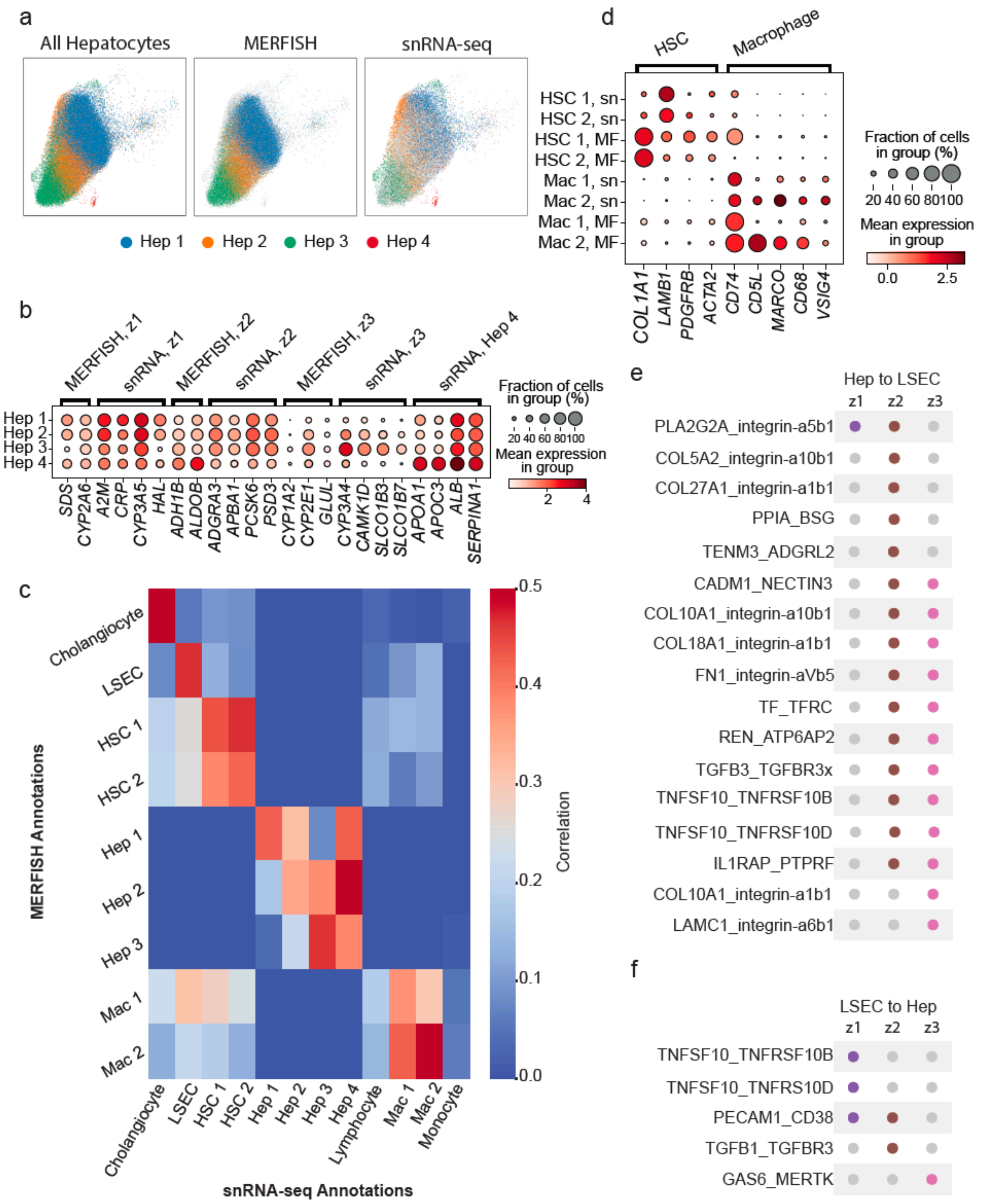
a
All Hepatocytes
MERFISH
snRNA-seq
Hep 1 Hep 2 Hep 3 Hep 4
b
MERFISH, z1
snRNA, z1
MERFISH, z2
snRNA, z2
MERFISH, z3
snRNA, z3
snRNA, Hep 4
Hep 1
Hep 2
Hep 3
Hep 4
SDS CYP2A6 A2M CRP CYP3A5 HAL ADH1B ALDOB ADGRA3 APBA1 PCSK6 PSD3 CYP1A2 CYP2E1 GLUL CYP3A4 CAMK1D SLCO1B3 SLCO1B7 APOA1 APOC3 ALB SERPINA1
Fraction of cells in group (%)
20 40 60 80100
Mean expression in group
2 4
c
MERFISH Annotations
Cholangiocyte
LSEC
HSC 1
HSC 2
Hep 1
Hep 2
Hep 3
Mac 1
Mac 2
Cholangiocyte LSEC HSC 1 HSC 2 Hep 1 Hep 2 Hep 3 Hep 4 Lymphocyte Mac 1 Mac 2 Monocyte
snRNA-seq Annotations
Correlation
0.5 0.4 0.3 0.2 0.1 0.0
d
HSC
Macrophage
HSC 1, sn
HSC 2, sn
HSC 1, MF
HSC 2, MF
Mac 1, sn
Mac 2, sn
Mac 1, MF
Mac 2, MF
COL1A1 LAMB1 PDGFRB ACTA2 CD74 CD5L MARCO CD68 VSIG4
Fraction of cells in group (%)
20 40 60 80100
Mean expression in group
0.0 2.5
e
Hep to LSEC
z1 z2 z3
PLA2G2A_integrin-a5b1
COL5A2_integrin-a10b1
COL27A1_integrin-a1b1
PPIA_BSG
TENM3_ADGRL2
CADM1_NECTIN3
COL10A1_integrin-a10b1
COL18A1_integrin-a1b1
FN1_integrin-aVb5
TF_TFRC
REN_ATP6AP2
TGFB3_TGFBR3x
TNFSF10_TNFRSF10B
TNFSF10_TNFRSF10D
IL1RAP_PTPRF
COL10A1_integrin-a1b1
LAMC1_integrin-a6b1
f
LSEC to Hep
z1 z2 z3
TNFSF10_TNFRSF10B
TNFSF10_TNFRS10D
PECAM1_CD38
TGFB1_TGFBR3
GAS6_MERTK

**Supplementary Figure 4. Integrating MERFISH with snRNA-seq. a.** UMAP of all hepatocyte cells from MERFISH and snRNA-seq (left), MERFISH alone (middle), and snRNA-seq alone (right). Colors indicate zone 1, 2, and 3 hepatocytes. A fourth cluster of hepatocytes (red) is only identified by snRNA-seq. For the MERFISH plot, cells from snRNA-seq are shown in gray, and for the snRNA-seq plot, cells from MERFISH are shown in gray. **b.** Dotplot of the expression of genes from snRNA-seq data by zone. Genes were either identified by MERFISH or snRNA-seq, as indicated. Expression of these genes is also shown for hepatocyte cluster 4, including additional genes enriched in this cluster. The size of the dot represents the percentage of cells expressing a specific RNA, and the color intensity indicates mean expression. **c.** Heatmap comparing correlation in expression of targeted MERFISH library between MERFISH (y-axis) and snRNA-seq (x-axis) clusters. Pearson correlation is indicated by color (far right) with the minimum and maximum set to 0.0 and 0.5, respectively. **d.** Dotplot showing expression of genes marking macrophages and HSCs in MERFISH (MF) and snRNA-seq (sn). The size of the dot represents the percentage of cells expressing a specific RNA, and the color intensity indicates mean expression. **e.** Receptor-ligand interactions were mapped between each zone of hepatocytes (zone (z) 1, z2, z3) and LSECs based on MERFISH. Only interactions that are uniquely enriched by hepatocyte zonation are shown. Receptor-ligand interactions characterized by ligand expression on hepatocytes and receptor expression in LSECs are shown. Gray circles indicate no interaction, and colored circles (zone 1 hepatocytes - purple; zone 2 hepatocytes - brown; zone 3 hepatocytes - pink) indicate receptor ligand co-expression. **f.** Receptor-ligand interactions between hepatocytes and LSECs characterized by receptor expression in hepatocytes and ligand expression in LSECs.

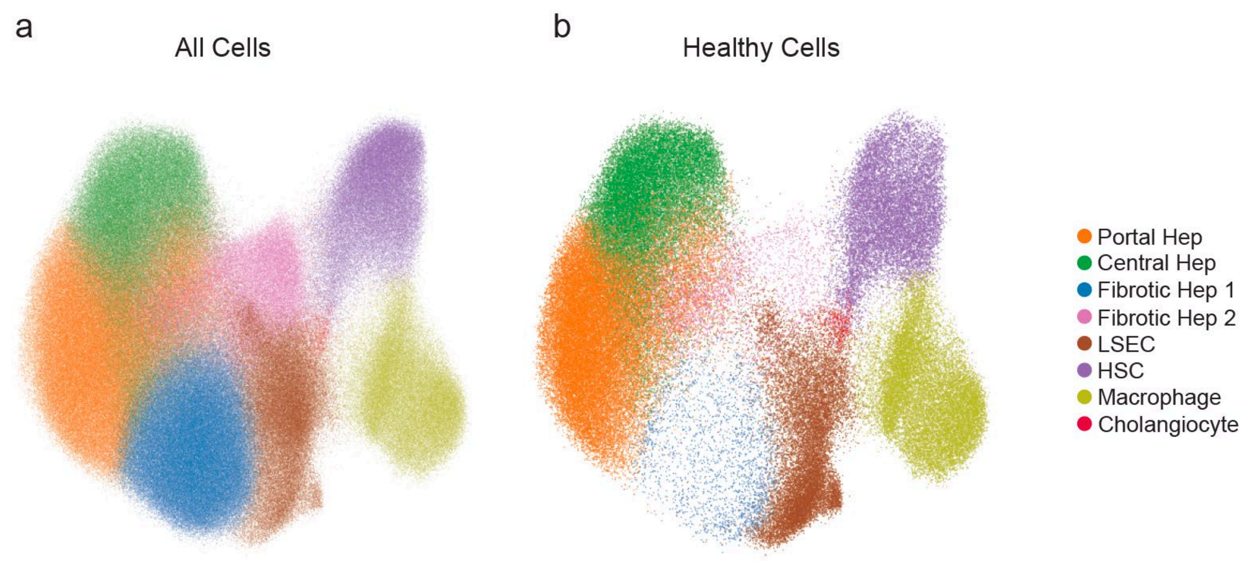


**Supplementary Figure 5. Merged healthy and fibrotic MERFISH data. a.** UMAP of all healthy and fibrotic cells measured with MERFISH. **b.** UMAP as in (**a**) but displaying only the cells imaged from healthy samples, highlighting the loss of Fibrotic Hep 1 (blue) and Fibrotic Hep 2 (pink) clusters in healthy samples.
